# Previous breeding success and carrion substrate together influence subsequent carrion choice by adult *Nicrophorus vespilloides*

**DOI:** 10.1101/2023.04.19.537504

**Authors:** Hyun Woo Park, Swastika Issar, Rebecca M. Kilner

## Abstract

Insects can adjust their behaviour in relation to experience in a wide range of contexts including foraging, mate selection, and choice of oviposition site. Here we investigate whether burying beetles modulate their choice of carrion in relation to the outcome of their past breeding experience. Burying beetles require small vertebrate carrion to reproduce. Beetle parents convert carrion into an edible nursery for their larvae, whom they typically care for throughout larval development. We tested whether a beetle’s past breeding experience influenced its subsequent choice of carrion, when presented simultaneously with either a dead mouse or a dead chick. We found that both male and female beetles favoured the same carrion as used in their first breeding attempt – but only if they had produced many larvae and only if they had previously bred on a mouse. Beetles that had produced fewer larvae on a dead mouse switched to favouring dead chicks in their second breeding attempt. Those that had bred on a dead chick chose carrion at random subsequently, regardless of their previous breeding success. Our general conclusion is that burying beetles can integrate different sources of information about their past breeding experience with current cues when selecting carrion for reproduction.

## Introduction

Choosing the right habitat and substrate for breeding is crucial for insects, especially when the available habitat conditions are variable. Holometabolous insects commonly produce larvae that are confined to the area surrounding the oviposition site until they gain mobility as adults. For example, many phytophagous insects remain on the same host plant until they reach adulthood (Tabashnik and Slansky, 1987), while parasitoid wasps develop and feed within a host species, before emerging to disperse away (Eggleton and Gaston, 1990). The parents’ choice of habitat for breeding can, therefore, strongly influence offspring survival and fitness (Akhtar and Isman, 2003).

One of the strategies used by insects to increase their chances of successfully reproducing is to evaluate the quality of the habitat by interacting with the habitat prior to oviposition (Jones and Agrawal, 2016). This has commonly been studied in phytophagous insects. For example, *Manduca sexta* adults lay more eggs on plants to which nectar has been added experimentally than on those without the added nectar, perhaps because this indicates the plant is of higher quality (Adler and Bronstein, 2004). Conversely, monarch butterflies (*Danaus plexippus*) lay fewer eggs on milkweed plants with higher cardenolide (a toxic defence compound) concentrations in the nectar because choosing plants with lower toxicity increases larval fitness (Jones and Agrawal, 2016).

An individual’s past experience with a certain habitat type could also be a useful source of information for evaluating habitat quality. This is observed in larvae in response to nutritional resources. Larvae of the spotted lady beetle *Coleomegilla maculata* (Coleoptera: Coccinellidae) learn about food quality, increasing the rate at which they reject poor quality food with experience (Boivin et al., 2010), while woolly bear caterpillars (Lepidoptera: Erebidae) avoid food that has previously caused illness (Dethier, 1980). In addition, it has been suggested that some species of phytophagous insects can increase the survival of their young by choosing to oviposit on host plant species that the adult successfully consumed as a larva (also known as the Hopkins’ host selection principle; (Hopkins, 1917). This has been studied in a number of species, but evidence for this idea is divisive (Akhtar and Isman, 2003; Chen et al., 2019; Chow et al., 2005; Craighead, 1921; Janz et al., 2009; Ning et al., 2018; Rausher, 1983; Trematerra et al., 2013). Similar claims have been made for parasitoids and preference for their host species, although it is uncertain whether host cues are picked up during the larval stage or in the early adult phase during eclosion (Emden et al., 1996; Gandolfi et al., 2003; Sasakawa and Kon, 2018).

Here we investigate whether burying beetles *Nicrophorus vespilloides* (Coleoptera: Silphidae) likewise integrate past experience with current cues in selecting the carrion they will breed upon. Burying beetles breed on small dead vertebrates such as mice or birds (Milne and Milne, 1976; Pukowski, 1933; Royle et al., 2013; Scott, 1998a). Adult burying beetles can locate carcasses to breed on from up to several kilometres away through olfaction, using antennal clubs with sensitive chemoreceptors (Abbott, 1927; Boeckh, 1962; Kalinová et al., 2009; Smith and Heese, 1995). When adults arrive on a carcass, they mate and prepare the carcass for reproduction by removing the fur or feathers, rolling it into a ball, and burying it (Scott, 1998a Milne and Milne, 1976). Females lay eggs in the soil surrounding the carcass during carcass preparation. When the larvae hatch, they enter the carcass and the parents feed and defend them (Pukowski, 1933; Sakaluk et al., 1998; Smiseth and Moore, 2002). About one week after mating, larval development is complete, and larvae disperse away from the carcass into the soil to pupate. Adults then search for fresh carrion and further breeding opportunities. It has been suggested that burying beetles breed more than once in the wild (Billman et al., 2014; Cotter et al., 2011; Creighton et al., 2009; Ward et al., 2009).

Burying beetles are known to adjust their behaviour in response to changing circumstances. For example, while rearing their larvae, burying beetles secondarily adjust their brood size based on the size of the carrion through filial cannibalism (Bartlett, 1987; Bartlett and Ashworth, 1988). In addition, Billman et al. (2014) showed that female burying beetles that previously bred on smaller carrion (which are therefore of lower quality) produced more offspring when they were given larger carrion to breed on next compared with beetles that previously bred on larger carrion. As a final example, adult females that experienced contests for carrion produced larger broods than beetles without prior experience of a contest (Pilakouta et al., 2016). Owing to these features, burying beetles provide an interesting subject for the study of plasticity in reproductive behaviour and offer a new ecological context for testing the generality of conclusions about egg-laying and substrate selection derived from other insect taxa.

To test the effect of past breeding experience on future carrion choice, we gave sexually mature beetles either a dead mouse carrion or a dead chick to breed upon and rear offspring. When larvae had completed development, the adult beetles were removed. One week later, the adults were presented simultaneously with a mouse carcass and a chick carcass and then allowed to choose one for reproduction. We tested whether the carcass-type they had used previously for reproduction influenced their subsequent choice of carrion, and whether this choice was modulated by their breeding success on the previous carrion substrate.

## Materials and Methods

### Laboratory Populations

The experimental populations originated from wild beetles collected from trapping sessions in Thetford Forest in August and September, 2017. The collected beetles were bred with each other in the laboratory, using mouse carrion. The population was organised into two blocks based on the time of trapping. The block descending from beetles caught in the first trapping session was named T1, while those from beetles caught in the second session gave rise to block T2. These populations were maintained as independent populations, without crossing in between populations. T1 and T2 blocks were offset by two weeks. Pairing siblings and cousins was avoided to prevent inbreeding depression. T1 and T2 populations were bred on mouse carrion at each generation using the husbandry techniques described below. Individuals were taken from the T1 and T2 populations at generation 20 for use in the experiment described here. No laboratory bred individuals were released back into the wild.

### General husbandry

Carrion was stored in freezers in the laboratory and left to thaw for 30-60 min at room temperature. Each mouse carcass was placed at the centre of a clear plastic box (17 cm × 12 cm × 6 cm), filled to 2 cm depth of moist Miracle-Gro compost, before adding a pair of beetles. Breeding boxes were placed in cupboards in the dark for eight days, by which point the final instar larvae were starting to disperse from the carrion into the compost. The larvae were removed, placed in gridded (5 × 5) boxes filled with fine compost, with one larvae in each cell, up to 25 per brood. The larvae were stored in these boxes for 17 days, during which they pupated and eclosed. Newly eclosed adult beetles were then placed individually in small plastic boxes (12 cm × 8 cm × 2 cm) containing moist compost and 0.5 grams of minced beef as food. The adult beetles were kept in these boxes for two weeks until they reached sexual maturity, at which they were paired and used in the next round of breeding.

## Experimental Methods

### Overview (see Figure 1a)

The experiment was divided into two steps. In step 1, we manipulated the beetles’ breeding experience by exposing the pair to one of two different types of carrion: either a dead mouse (of the same type that they had been raised on as larvae) or a dead domestic chick (unfamiliar carrion). We measured the pair’s reproductive success at the point when larvae dispersed away into the soil to pupate and then split up the pair. In step 2, we presented these adults with a simultaneous choice of carrion: either a dead mouse or a dead domestic chick. Males and females were tested individually. We recorded which carrion the beetles prepared for reproduction and compared their choice of carrion to their previous reproductive success.

**Figure 1.**
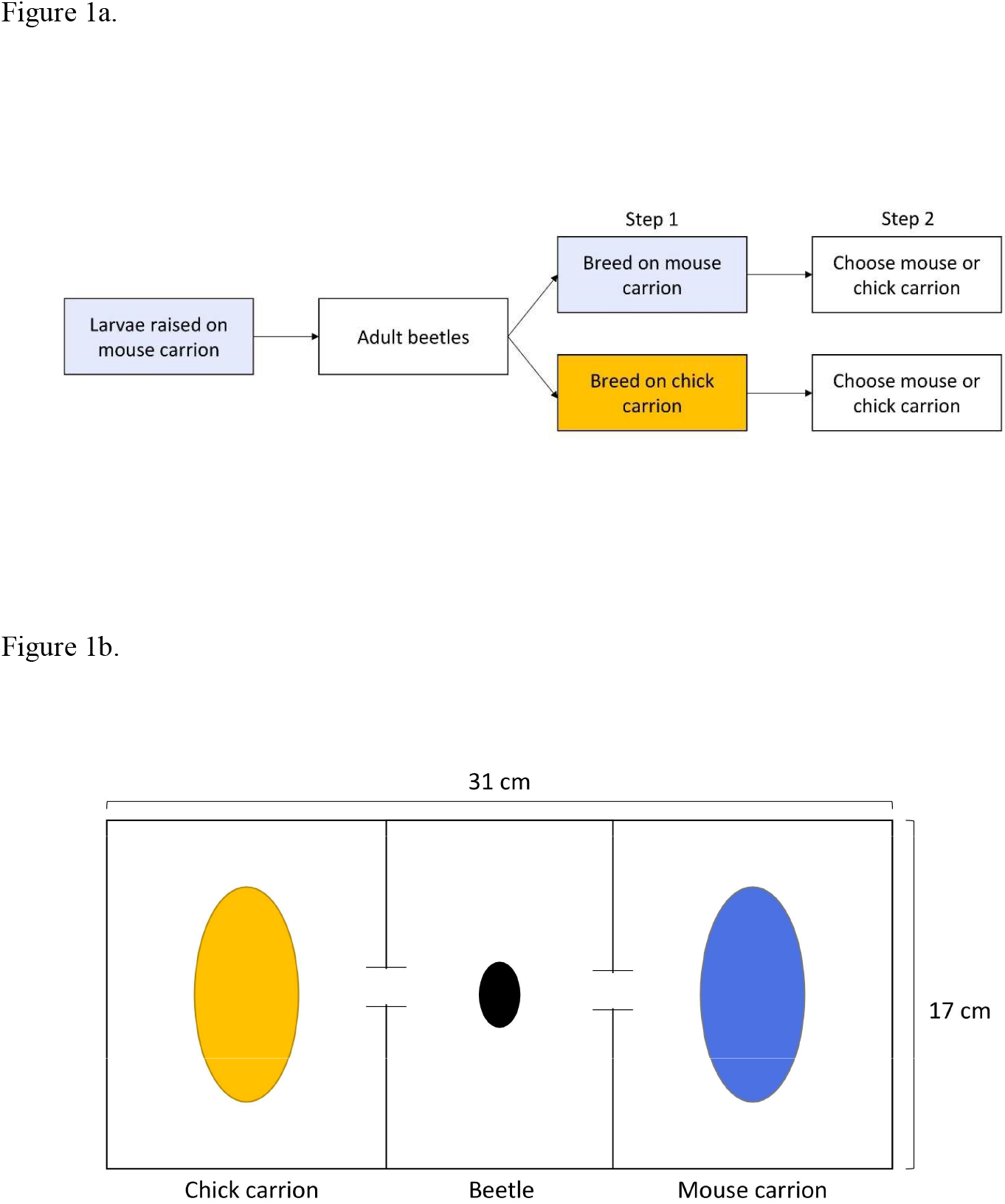
a) Overview of experimental design and b) schematic plan view of the chamber used for the preference tests in Step 2. The chamber was lined with compost, and one dead mouse and one dead chick placed on top of the compost on each side (the location of the carrion type was alternated between trials). The openings in the dividers allowed a beetle placed in the centre to move freely before settling on and processing the carrion of their choice. Carrion mass was controlled to fall between 19-25 g for both carrion types.

### Detailed Methods

#### Step 1

Two weeks post eclosion, when the adult beetles reached sexual maturity in generation 20, they were paired with individuals from the same replicate populations (i.e. T1 individuals were paired together and T2 individuals were paired together). The size of each beetle was recorded by measuring pronotum width using Vernier callipers. Each pair was provided with either a mouse carcass or a chick carcass to breed on, as described above. Carrion mass was controlled to fall between 19-25 g for both carrion types. Eight days after pairing, when the larvae were dispersing away from the carrion, larval density was measured as the total number of larvae in a brood divided by carrion mass (measured at pairing). The parent beetles were removed and kept individually for one week in small plastic boxes, filled with compost and fed a piece of minced meat.

#### Step 2

One week after breeding, adults were tested for their carrion preference. A total of 151 beetles were chosen at random for testing: 72 beetles from T1 block (36 bred on mouse carrion (18 males and 18 females), 36 bred on chick carrion (18 males and 18 females) were tested. 80 beetles from T2 block (40 bred on mouse carrion (20 males and 20 females), 40 bred on chick carrion (20 males and 20 females)). Preference tests were conducted using custom-built chambers, each made from a transparent plastic box (31 cm × 17 cm × 10 cm) (Figure 2). Each box was filled with a 3 cm layer of Miracle-Gro compost and divided into three chambers using two dividers, each made with plastic with a small opening (2 cm across × 4 cm height) in the middle (Figure 1b). This allowed the beetles to move in between chambers. Mouse and chick carrion were thawed for 72h before the experiment. A dead mouse was placed at one end of the box, while a dead chick placed at the other end. Carrion mass was controlled to fall between 19-25 g for both carrion types. The experiment began when a test beetle was placed in the central compartment. The choice chamber was then sealed and kept in a dark room to simulate a nocturnal setting. 30 h after the start of the experiment, the choice chamber was opened and the carrion that had been chosen and processed by beetles was recorded. Signs of processing included burying, removal of fur/feathers and punctures in the skin. Beetles that processed both (10 beetles across all populations) or neither type of carrion (5 beetles across all populations) were marked as such and removed from the analysis. After each test session, the boxes and dividers were thoroughly washed and cleaned to remove any potential trace of the carrion or beetles from previous tests before use in the next test session.

**Figure 2.**
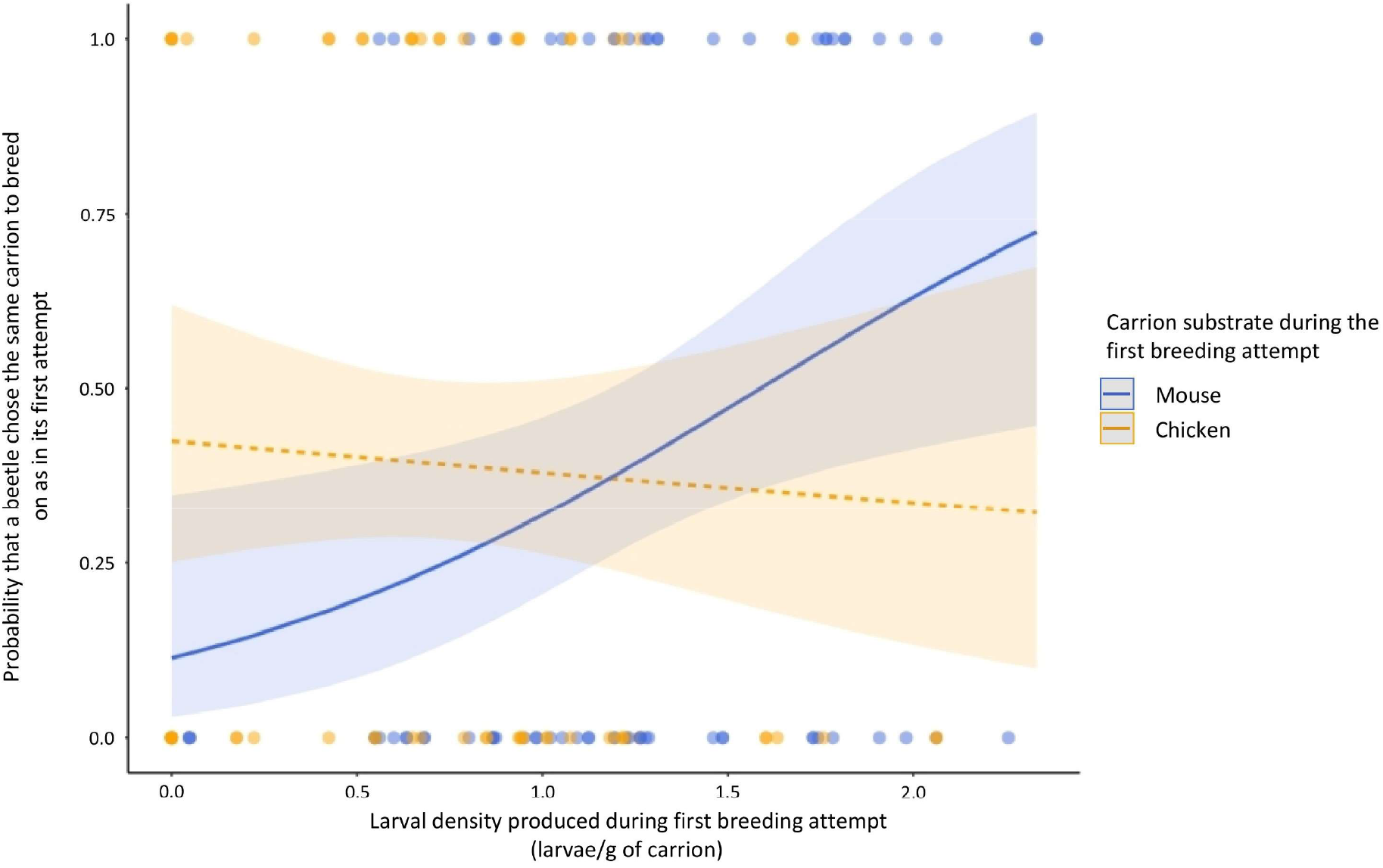
Carrion choice in relation to the carrion type on which the beetle bred, and the larval density they produced, in Step 1 of the experiment. Predicted values are shown for beetles that bred on mouse carrion (in blue) and on chick carrion (in yellow), together with 95% confidence intervals. Each datapoint represents an individual test beetle (N = 137 beetles).

### Statistics

All statistical analyses were performed using RStudio (version 4.2.1). Carrion choice was analysed using logistic mixed effects model, and scored as either 1 (chose same carrion as the one it bred on in step 1) or 0 (chose different carrion from the one it bred on in step 1). We included as variates the carrion type the beetle had previously bred on, its sex, size, density of larvae produced during the first breeding attempt, and the interaction between carrion type and larval density. Correlation between variates was tested using variance of inflation factors (VIFs), using a cut-off point of 5 (Sheather, 2009). Non-significant interaction terms and explanatory variables were removed from the model using backward stepwise elimination.

## Results

The carrion type that the beetles were bred on in Step 1 influenced their subsequent carrion preference, through an interaction with their previous breeding success (Table 1, GLM followed by type III Wald chi square tests, p<0.05).

**Table 1.** Results from logistic regression to explain variation in carrion preference. P values were calculated based on type III Wald chi square tests. Number of observations was 137, and the degrees of freedom of the model was 133.

| Variates and factors | Estimate | SE | z value | p |
| --- | --- | --- | --- | --- |
| Intercept | -0.3038 | 0.4031 | -0.754 | 0.4511 |
| Larval density | -0.1886 | 0.4392 | -0.429 | 0.6676 |
| **Carrion type bred on in Step 1 [Mouse]** | -1.7507 | 0.8291 | -2.111 | **0.0347\*** |
| **Larval density: Carrion type bred on in Step 1 [Mouse]** | 1.4815 | 0.6817 | 2.173 | **0.0298\*** |
| Non-significant variables (eliminated step-wise from the model) |  |  |  |  |
| Beetle size | -0.4240 | 0.4886 | -0.868 | 0.3855 |
| Population [T2] | -0.33661 | 0.39386 | -0.855 | 0.3927 |
| Sex [Male] | 0.3170 | 0.3708 | 0.855 | 0.3925 |

For beetles that were bred on mice carrion in Step 1, the preference for a mouse carcass in Step 2 increased with their previous breeding success. Beetles that had produced larger broods on mouse carrion were more likely to subsequently prefer mice carrion compared to beetles that had produced smaller broods on mice in their first breeding attempt (Table 1, Fig. 2). The same was not true for beetles that previously bred on chicks (Table 1, Fig. 2). Population block, beetle size (width of pronotum), and sex did not significantly explain variation in carrion preference.

## Discussion

In this study, we found that previous breeding experience affected the subsequent choice of carrion by adult burying beetles. Beetles that bred initially on mice carrion and that produced high larval densities were more likely to choose mice carrion subsequently, whereas those that produced low density broods on mice carrion were less likely to choose mice subsequently and more likely to opt for the alternative resource (chick carrion). On the other hand, beetles that were bred on chick carrion showed no particular preference or either carrion type, regardless of their previous breeding success on the chick carrion.

Multiple factors could have influenced carrion choice. An important factor could be difference in the quality of mouse and chick carrion as breeding options. In this study (Fig. 2) and other work (Issar et al., unpublished data) using laboratory populations of burying beetles, the mouse carrion has been reported to be a higher quality resource, yielding greater brood size and larval density compared to chick carrion. The reason for this is unknown; although burying beetles are known to use small bird carrion for breeding, the chick carrion used in this study was commercially produced and may be nutritionally different from wild birds.

However, carrion preference was not explained by differences in carrion quality alone. If carrion choice was based on carrion quality, or an innate preference hierarchy as observed in other insects (Scriber, 1993; Trematerra et al., 2013), then beetles from both treatments would be expected to show higher preference for mice carrion. Likewise, carrion preference was not explained either by previous breeding experience alone or by beetle size alone. The switch only happened if beetles had previously produced a brood with low density of larvae and on mouse carrion.

These findings suggest that beetles are using cues from both the previous carrion type and their reproductive success on the carcass. This implies that beetles can identify the type of carrion they used previously, and also assess their own breeding success. Information about the carrion type is most likely to be gained from olfactory or gustatory cues obtained while preparing and consuming the carrion (Abbott, 1927; Boeckh, 1962; Kalinová et al., 2009). Larval density could be obtained directly since adult burying beetles remain on the carcass to provide care for their larvae after hatching (Milne and Milne, 1976; Pukowski, 1933; Royle et al., 2013; Scott, 1998b; Smiseth and Moore, 2002). In this process, burying beetles directly interact with their own larvae (Smiseth and Parker, 2008), potentially giving them the opportunity to evaluate how many offspring they have produced.

Rejecting a resource on which beetles had not been very successful and seeking a novel resource could be an adaptive strategy that increases success rate in the next breeding round. Similarly, the tendency for burying beetles to show increased preference for previously unknown resource after a poor breeding experience is consistent with how many animals react to an unsuccessful reproductive attempt in their next opportunity. For example, males of the eastern amber-wing dragonfly *Perithemis tenera* (Odonata: Libellulidae) hover around ponds wherein females mate and oviposit. Field experiments have shown that males that have failed to mate were more likely to move to different ponds the following day to seek new opportunities (Switzer, 1997a, 1997b). Similar results have been reported from dispersal patterns of birds (Beletsky and Orians, 1991; Danchin et al., 1998; Newton and Marquiss, 1982). For example, collared flycatchers (*Ficedula albicollis*) that were unsuccessful at mating or breeding were more likely to disperse further to new territories than successful individuals (Doligez et al., 1999; Pärt and Gustafsson, 1989). Bobolink (*Dolichonyx oryzivorus*) that had failed to raise their young on low quality sites were much less likely to return to the same nesting sites the next year, while successful individuals returned to the same sites at equal rates, regardless of habitat quality (Bollinger and Gavin, 1989).

Interestingly, past breeding experience only affected carrion choice for beetles that bred on mouse carrion. For beetles that were given a chick carrion to breed in stage 1, the larval density did not significantly affect carrion choice. Instead, these beetles showed a slight preference for mouse carrion that was not statistically different from random choice, regardless of larval density produced. A possible reason is that chick carrion is a lower quality resource than mouse carrion, and this may have overshadowed any effect of brood size. Poorly fed individuals tend to be less choosy (Stamps, 2006). Context-dependent induction of preference for habitat has been demonstrated in phytophagous insects. For example, female Egyptian cotton leafworms (*Spodoptera littoralis*) were exposed to olfactory cues from a host plant while being fed on high or low-quality artificial diets. They showed increased oviposition preference for the host plant only when they were fed on a high-quality diet (Lhomme et al., 2018).

In conclusion, this study suggests that the carrion preference of burying beetles can be affected by their past breeding experience when a mouse carrion was used initially for breeding. The challenge for future work is to identify the cues that underpin this behavioural switch.

